# Evaluation of Neuropharmacological Effects of Different Chemical Extracts of *Flemingia Stricta* (Roxb.) Leaves

**DOI:** 10.1101/2020.04.09.034553

**Authors:** Md. Shahrear Biozid, Mohammad Nazmul Alam, Md. Jainul Abeden, Md. Masudur Rahman, Md. Rafikul Islam

## Abstract

**Background:** Traditional preparation of the leaf of *Flemingia stricta* (Fabaceae) Roxb., a medicinal plant of the Indian subcontinent, has been used for treatment of different diseases as herbal preparation. Our purpose was to analyze Neuropharmacological effects of different chemical extracts of *Flemingia stricta* Roxb. as particular form of behavioral inhibition that occurs in response to novel environmental events.

**Methods:** In present study, the anxiogenic activity of crude extracts of *Flemingia stricta* leaves was determined using standard animal behavioral models, such as hole cross and open field; Sedative property and anxiolytic potential were assessed by conducting thiopental sodium induced sleeping times tests and elevated plus maze test respectively.

**Results:** The crude extracts at both dose (200 and 400mg/kg) exhibited a significant (P<0.05, P<0.01) dose-dependent suppression of motor activity and exploratory activity of mice in both open field and hole cross test. In anxiolytic study, extracts displayed increased percentage of entry into open arm at the dose of 200 and 400mg/kg. Extracts produced a significant (P<0.05, P<0.01) increase in sleeping duration and reduction of onset of sleep compared to sodium thiopental at both doses (200 and 400mg/kg).

**Conclusion:** This study demonstrates that the treated extracts have significant central nervous system depressant effect. Further studies on the active constituent of the extract can provide approaches for therapeutic intervention.

## Background

Presently herbal drugs are wide-spoken as green medicine for their safe and trustworthy health care paradigms. Traditional herbal medicines a rising interest since a couple of decades due to their incredible pharmacological activities, economic viability and less side effects in different healthcare managements rather than other systems [1]. Anxiety and depression are the most common psychiatric disorders; already cover 20% of the adult population, suffering from these illnesses at some time during their lives [2-4]. It has grown up to be an important area of research interest in psychopharmacology during this decade [5].

Drugs acting on the central nervous system (CNS) are still the most widely used pharmacological agents [6]. Benzodiazepines are among the most prescribed and effective anti-anxiety drugs used worldwide [7]. Barbiturates and Ethanol are also frequently used. Both barbiturates and benzodiazepines show their CNS effect by interaction with postsynaptic gamma aminobutyric acid receptor (GABA_A_ receptor) [8]. The most serious shortcoming of barbiturates as a depressant is linked to their narrow margin of safety, and only 10 times of their therapeutic dose may be lethal [9]. Moreover barbiturates can grow both psychological and physiological dependence [10, 11]. Benzodiazepines are the most commonly used CNS depressant which lead to tolerance and physical dependence, for example diazepam typically produces sedation at dose of 5 to 10 mg in user of first time, but those who repeatedly use it may become tolerant to doses of several hundred milligrams [12]. Ethanol produces its depressant action by changing membrane fluidity and interaction with the GABA system [9, 13]; also it has a tolerance and physical dependence activity. Statistically it has been showed; alcohol addiction in American society is 5% to 10% for men and 3% to 5% for women [14]. A natural CNS depressant with minimum or no toxicity is therefore, essential.

*Flemingia stricta* (Fabaceae) Roxb. is an erect subshrub, distributed in the Southeast Asian country such as-Bangladesh, Bhutan, China, India, Indonesia, Laos, Myanmar, Philippines, Thailand and Vietnam [15, 16]. In Bangladesh, it is available in Chittagong, Chittagong Hill Tracts and Sylhet. It is known as Charchara (in Bangla) and Krangdunaduepay, Sai Kheu (Marma); Keramkana(Tripura); Tamatamaking (Khumi) and Harsanga, Khaskura, Uskura (Chakma) in local tribes of Chittagong, Bangladesh[17]. *Flemingia stricta* is used by Chakma healers for treatment of polio. The species is also used to treat rheumatism followed by bone fracture, cough, asthma, goiter, urinary problems, snake bite, insect bite, leprosy, tumor and cancer, caries, hysteria, tuberculosis, insomnia and intestinal worms [18-20].

According to the literature review on this plant, it showed that the plant has been used as traditional medicine for the treatment of various diseases for many years. Therefore, we undertook the study to assess the neuropharmacological potential of *F. stricta* leaves, by using different animal models and studying the effect of the different chemical plant extracts on their exploratory behavior.

## Methods

### Plant collection and identification

Whole plants of *F. stricta* were collected from Bhatiary, Chittagong region, Bangladesh. The plants were identified by Dr. Shaikh Bokhtear Uddin, Taxonomist and Professor, Department of Botany, University of Chittagong, Chittagong, Bangladesh.

### Preparation and extraction of leaf extract

The collected leaves were thoroughly washed with distilled water and dried under the shade. The dried sample was coarsely powdered (500 g) and extracted with methanol for 3 days to allow total extraction process. After that the plant extract was filtered with sterilized cotton filter and the filtrate was gathered in a beaker. The plant extracts then kept in a water bath at 60 °C to evaporate the solvent from the solution. The container allowed to airtight for 72 h and filtrate thus obtained was concentrated by using a rotary evaporator. The extract was divided into two portions. One portion (2.5 g) was poured into glass vials to be tested as crude methanol extract, whereas the second portion (8 g) was dissolved in concentrated methanol and partitioned successively into four different extractives [21]. The fractions were then concentrated using a rotary evaporator which results ethyl acetate fraction (yield weight 1.50 g), chloroform fraction (yield weight 1.25 g), petroleum ether fraction (yield weight 2.15 g) and aqueous fraction (2.75g).

### Animal

Male Swiss albino mice, 3-4 weeks old, weighing between 20-25g, were collected from the International Center for Diarrheal Disease and Research, Bangladesh. Animals were maintained under standard environmental conditions [temperature: (24±1) °C, relative humidity: 55%-65% and 12 h light/12 h dark cycle] and had free access to feed and water ad libitum. Prior to experimentation, animals were familiarized in laboratory condition for one week.

### Acute toxicity study

Mice were divided into control and test groups (n = 5). The test groups received the extract per orally at the doses of 600, 800 and 1000, mg/kg. Then the animals were kept in separate cages and were allowed to food and ad libitum. The animals were observed for possible behavioral changes, allergic reactions and mortality for the next 72 h [22].

### Neuropharmacological tests

The study was done to find out if extracts had any effect on the central nervous system. Effect on exploratory behavior of mice was evaluated by hole cross test and open field test. Elevated plus maze test was conducted for determination of anxiolytic activity whereas thiopental sodium induced sleeping time test was for sedative activity.

### Open field test

The method was adopted as described by Gupta et al [22]. In the open field test, the animals were divided into control, positive control and test groups containing 5 mice each. The test groups received extract of *F. stricta* at the doses of 200 and 400 mg/kg body weight orally whereas the control group received the vehicle (1% Tween 80 in water). The floor of half square meter open field was divided into a series of squares each alternatively colored black and white. The apparatus had a 40cm height walls. The number of squares traveled by the animals was counted for 5 min at 0, 30, 60, 90, 120 min after oral administration of both doses of the extract.

### Hole cross test

The apparatus was a cage of 30cm×20cm×14cm with a steel partition fixed in the middle, dividing the cage into two chambers. A hole of 3.5cm diameters was made at a height of 7.5cm in the center of the cage. Animals were randomly divided into control, positive control and test groups containing 5 mice each. The test groups were treated with extract of *F. stricta* at the doses of 200 and 400 mg/kg body weight orally whereas the positive control group with diazepam (1 mg/kg) and control group with vehicle (1% Tween 80 in water). Number of passages of the animals through the hole from one chamber to the other was counted for 5 min at 0, 30, 60, 90 and 120 min after oral administration of the extract as well as diazepam and vehicle [23]. The apparatus was thoroughly cleaned after each trial.

### Thiopental sodium induced sleeping time test

For the experiment, the animals were randomly allocated to four groups, each with 5 mice. The test groups were given the leaf extract of *F. stricta* at doses of 200 and 400 mg/kg body weight, while the positive control was treated with diazepam (1 mg/kg) and control group with vehicle (1% Tween 80 in water). Thirty minutes later, thiopental sodium (40mg/kg) was administered to each mouse to induce sleep. The animals were observed by placing them on separate chambers for the latent period (time between thiopental administrations to loss of righting reflex) and duration of sleep i.e. time between the loss and recovery of righting reflex. The onset of sleep and total sleeping time were recorded for control, positive control and test groups [24].

### Elevated plus maze test

The method initially suggested by Handley and Mithani were employed with minor modifications [25]. The apparatus consists of two open arms (5 × 10) cm and two closed arms (5 × 10 × l5) cm radiating from a platform (5 × 5) cm to form a plus-sign figure. The apparatus was situated 40cm above the floor. The open arms edges were 0.5cm in height to keep the mice from falling and the closed-arms edges were 15cm in height. Sixty minutes after administration of the test drugs, each animal was individually placed in the center of the EPM and was allowed 5 min for free exploration. Next, the number of open and enclosed arm entries, and time spent on open arms was manually registered [26]. Entry into an arm was defined as the point when the animal placed all four paws onto the arm. The percentage of open arm entries (100 × open/total entries) and the percentage of time spent in the open arms (100 × open/ (open + enclosed)) were calculated for each animal. Observations made from an adjacent corner produced significant (*p*< 0.05, *p* < 0.01) decreases of locomotion from its initial value during the period of the experiment (Table 1). Maximum suppression of locomotor activity was displayed at the dose of 400 mg/kg body weight, which was comparable to the reference drug diazepam.

**Table 1:**
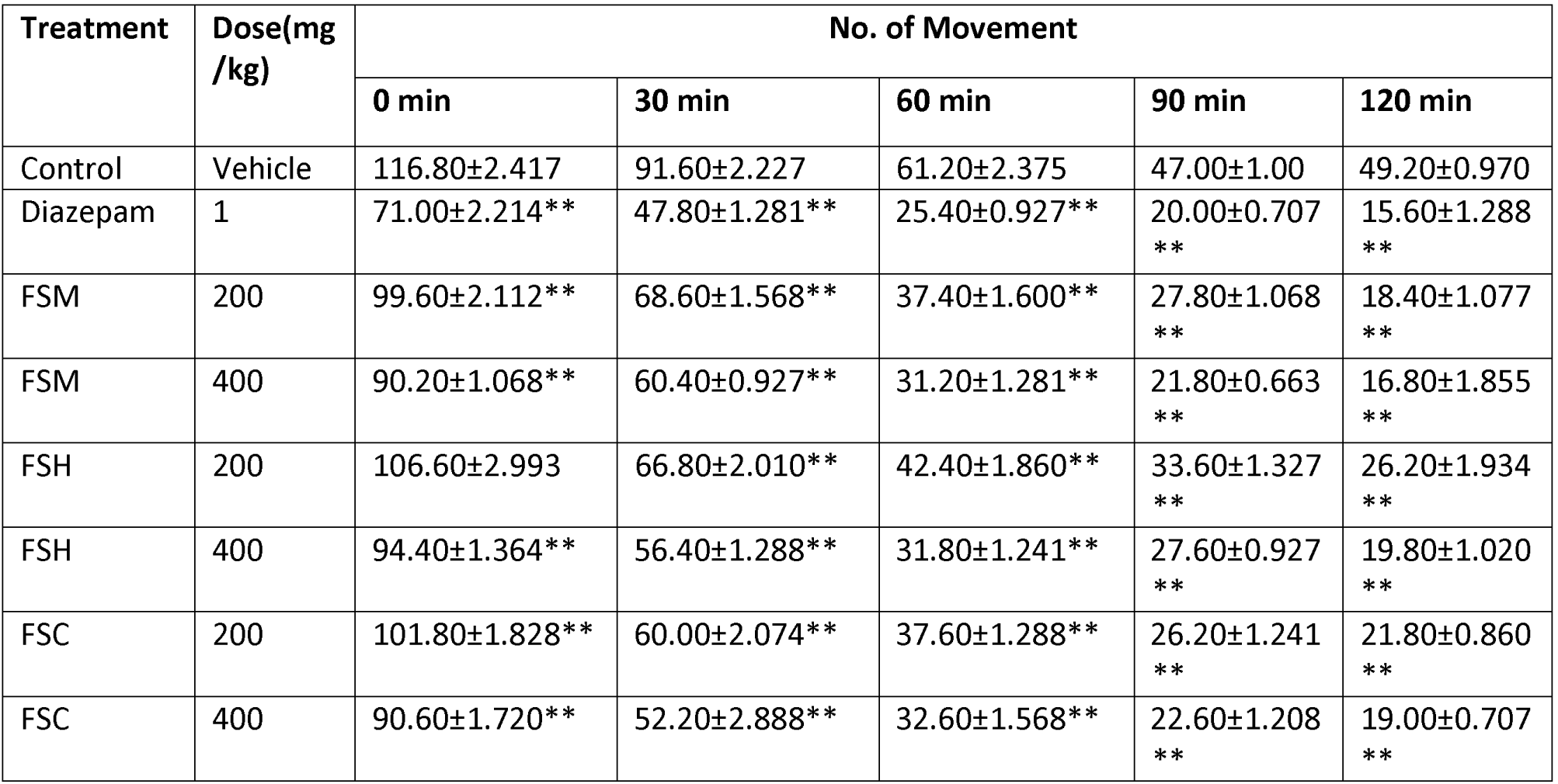
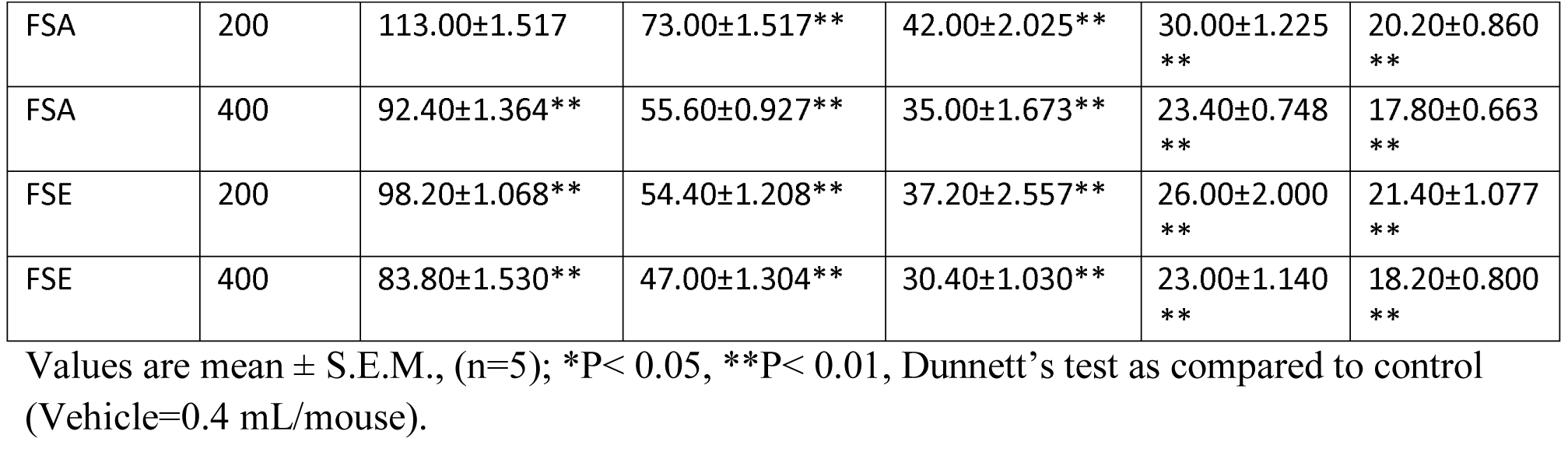
Effect of extracts of *F. stricta* on exploratory behavior on mice. (Open field test)

### Statistical analysis

The data was expressed as mean ± standard error of mean (S.E.M.). Statistical comparisons were performed using one-way ANOVA followed by post-hoc Dunnett’s test with the SPSS program (SPSS 20.0, USA). The values obtained were compared with the vehicle control group and were considered statistically significant when *P< 0.05, **P< 0.01.

## Results

### Neuropharmacological tests

#### Open field test

Open field test of *F. stricta* treated groups (200 and 400 mg/kg body weight) showed significant and dose-dependent reduction of movement from its initial value at 0 to 120 min (Table 1). The number of squares traveled by the mice was reduced significantly from its initial value at 0 to 120 min at the dose level of 200mg/kg and 400 mg/kg body weight (P<0.05) of the extracts from the leaves of *F. stricta* (Table 1).

#### Hole cross test

Hole cross test of *F. stricta* treated groups showed a decrease of movement from its initial value at 0 to 120 min. Results showed in the table (Table 2) suggests that the number of hole crossed from one chamber to another by mice was decreased significantly for all chemical extracts of *F. stricta* leaves compared to the standard drug Diazepam. But, at doses of 400 mg/kg (P<0.05), maximum suppression of locomotor activity was displayed which was comparable to the reference drug diazepam (Table 2).

**Table 2:**
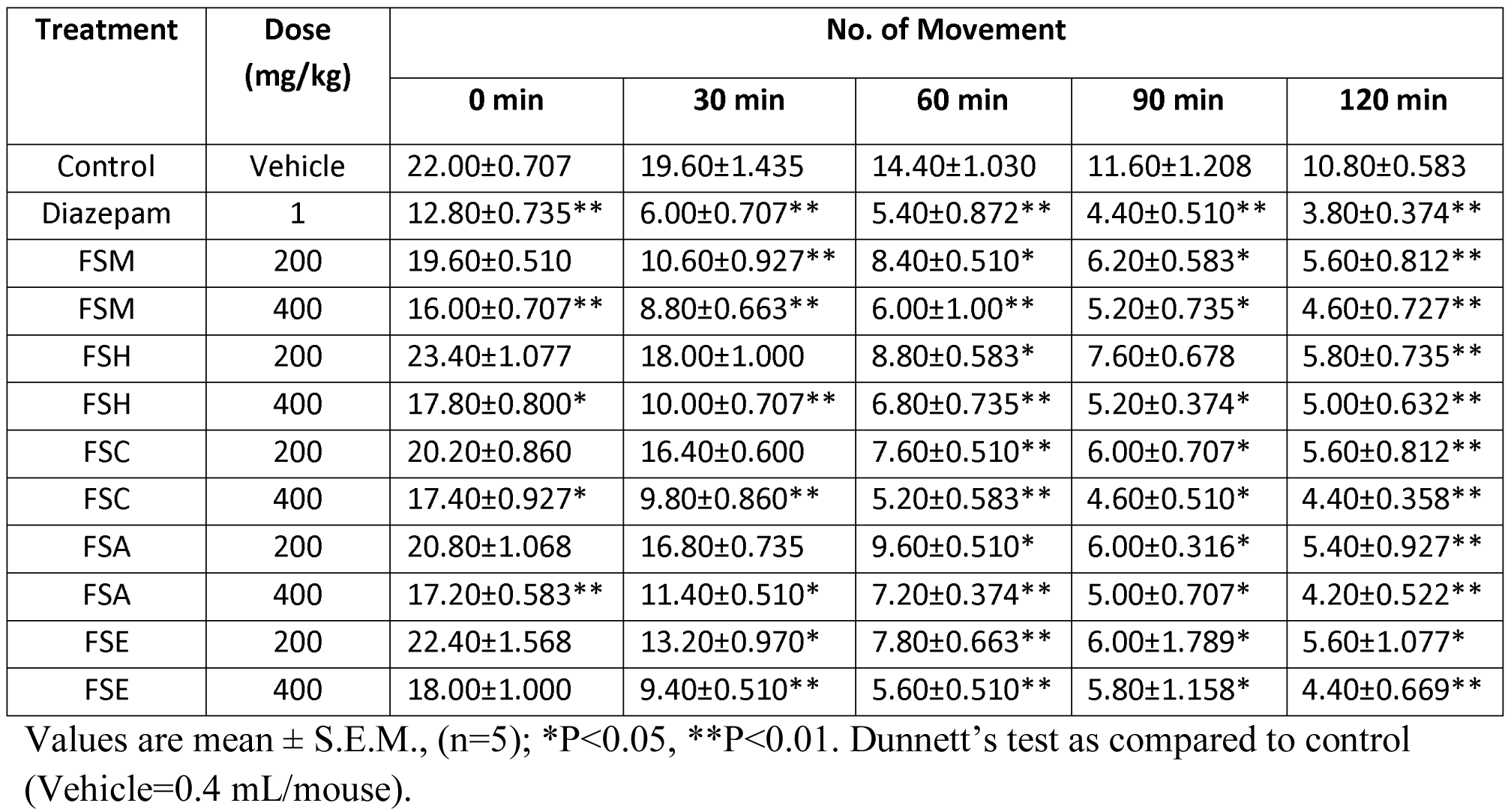
Effect of extracts of *F. stricta* on exploratory behavior on mice. (Hole cross test)

#### Thiopental sodium induced sleeping time test

In the thiopental induced hypnosis test, the extract at doses, 200 and 400 mg/kg showed a significant reduction in the time of onset of sleep in a dose-dependent manner; mostly in the case of methanol and ethanol extract of *F. stricta* leaves (Table 3). The effect of the extract (200 and 400 mg/kg) on the onset of sleep was comparable to that of standard drug Diazepam. Both doses of the extract potentiated the duration of thiopental sodium induced sleeping time in test animals compared to controls (Table 3).

**Table 3:** Effect of extracts of *F. stricta* on thiopental sodium induced sleeping time.

| Treatment | Dose(mg/kg) | Onset of sleep(min) | Duration of sleep(min) |
| --- | --- | --- | --- |
| Control | Vehicle | 42.50±1.059 | 47.70±0.574 |
| Diazepam | 1 | 13.88±1.151** | 142.14±3.075** |
| FSM | 200 | 13.70±1.537** | 97.86±4.853** |
| FSM | 400 | 7.66±0.731** | 119.00±4.226** |
| FSH | 200 | 16.98±0.664** | 85.78±3.229** |
| FSH | 400 | 10.64±0.670** | 101.06±1.997** |
| FSC | 200 | 15.84±0.828** | 89.26±2.839** |
| FSC | 400 | 10.04±0.451** | 103.06±1.492** |
| FSA | 200 | 15.34±1.332** | 109.00±3.730** |
| FSA | 400 | 11.98±0.828** | 126.42±3.441** |
| FSE | 200 | 13.16±0.832** | 112.20±2.663** |
| FSE | 400 | 5.66±0.510** | 126.20±3.097** |

Values are expressed as mean ± S.E.M, (n=4-5); \*P<0.05, \*\* P<0.01, Dunnett's test as compared to control (Vehicle=0.4 mL/mouse).

#### Elevated plus maze test

In the EPM, the behavior of mice model, as observed, confirmed the anxiolytic activity of diazepam as reported previously. The extracts of *F. stricta* at the dose of 400 mg/kg (P<0.01), significantly increased the percentage of entries of mice into the open arms, and the percentage of time spent in the open arms of the EPM as shown in Table 4. The effects of treatment of mice at the dose of 200 mg/kg on open arm entries and time spent in open arms were dose dependent. The number of closed arm entries and time spent in the closed arms was decreased significantly in the extract treated groups which were comparable with the standard diazepam.

**Table 4:** Effect of extracts of *F. stricta* on EPM test during 5 min test session.

| Treatment | Dose(mg/kg) | % Entry into open arm | % time spent in open arm |
| --- | --- | --- | --- |
| Control | Vehicle | 35.69±1.244 | 35.02±1.181 |
| Diazepam | 1 | 71.98±1.090** | 73.29±1.457** |
| FSM | 200 | 43.87±1.215** | 46.19±1.217** |
| FSM | 400 | 58.12±2.037** | 58.65±2.252** |
| FSH | 200 | 38.40±1.085 | 37.95±0.914 |
| FSH | 400 | 56.09±0.577** | 55.74±0.794** |
| FSC | 200 | 36.91±1.493 | 40.05±0.609* |
| FSC | 400 | 58.48±1.886** | 58.24±1.130** |
| FSA | 200 | 42.35±1.030* | 43.11±0.558** |
| FSA | 400 | 60.76±1.324** | 61.49±1.230** |
| FSE | 200 | 42.25±1.446* | 44.41±0.442** |
| FSE | 400 | 62.84±0.855** | 61.96±2.346** |

Values are expressed as mean ± S.E.M., (n=5); \*P<0.05, \*\*P<0.01, Dunnett's test as compared to control (Vehicle=0.4 mL/mouse).

## Discussion

The result of open field and hole cross tests showed that the studied plant decreased the frequency as well as the bountifulness of movements. Since the level of excitability of the CNS is measured by locomotor activity, this reduction in spontaneous motor activity that could be considered as the sedative effect of the plant extracts. Locomotor activity lowering effect was evident at the 2nd observation (30 min) and continued up to the 5^th^ observation period (120 min). So in addition, the study on locomotor activity, as measured by hole cross and open field tests, it can be stated that both doses of chemical extracts of *F. stricta* leaves decreased the frequency and the amplitude of movements. The above result also showed that crude extracts of *F. stricta* plant had strong sedative and hypnotic action that principally mediated in the CNS by the GABA_A_ receptor complex. Thiopental, a barbiturate drug, produce sedative-hypnotic result at a certain dose due to their interaction with GABA_A_ receptors which enhance the GABAergic transmission. It potentiates GABA activity, entering chloride into the neuron by prolonging the duration of chloride channel opening. On the other hand, thiopental can inhibit excitatory glutamate receptors. All of these molecular action lead to decrease of neuronal activity that supports the following reference substances which possess sedative action.

Substances which possess CNS depressant activity either decrease the time for the onset of sleep or prolong the duration of sleep or both [27, 28]. The method employed for this assay is considered as a very sensitive way to detect agents with CNS depressant activity [29]. The sedative effect recorded here may be linked to an interaction with benzodiazepines and related compounds that bind to receptors in the CNS and have already been identified in certain plant extracts. So the plant extract of *F. stricta* may contain alkaloids, glycosides, cardiac-glycosides, flavonoids, steroids, tannins, anthraquinone glycosides and saponins. The EPM is one of the most widely validated tests when the test drug increases open arms entries without altering the total number of arm entries and is highly sensitive to the influence of both anxiolytic and anxiogenic drugs acting at the gamma aminobutyric acid type A (GABAA) - benzodiazepine complex [30]. In EPM, normal mice will generally prefer to spend much of their allotted time in the closed arms. This preference appears to reflect an aversion towards open arms that are generated by the fears of the open spaces. Drug like diazepam that increases open arm exploration is considered as anxiolytic and the reverse holds true for anxiogenics [31]. In this study, we observed that the administration of different doses (200 and 400 mg/ kg body weight) of extracts of *F. stricta* induced an anxiolytic-like effect in mice, as it increased open arm entries and the time spent in the open arms of the EPM when compared to the control animals.

## Conclusion

All of the results were dose-dependent and statistically significant. Analyzing the results of the present study, it can be inferred that the crude extracts of *F. stricta* possess significant neuropharmacological activity. Therefore, we can suggest that the extract may fulfill the therapeutic need for the treatment of anxiety and related neuropsychiatric disorders. However, further investigation is necessary to determine the exact phyto-constituents and mechanism of action that are responsible for the biological activities of the leaf extracts of *F. stricta.*

## Abbreviations

F. stricta: *Flemingia stricta*;
EPM: Elevated plus maze;
GABA: Gamma-aminobutyric acid;
CNS: Central nervous System,
FSM: *F. stricta* Methanol,
FSH: *F. stricta* N-hexane,
FSC: *F. stricta* Chloroform,
FSA: *F. stricta* Aqueous,
FSE: *F. stricta* Ethanol.

## Ethical approval

The set of rules followed for animal experiment were approved by the institutional animal ethics committee, Department of Pharmacy, International Islamic University Chittagong, Bangladesh according to governmental guidelines. Pharmacy P&D 59/04-10.

## Consent for publication

All authors read and approved the final content of this manuscript for publication.

## Authors’ contributions

MSB, MNA and MMR proposed and designed the study. MSB, MNA and MJB conducted all laboratory experiments; MNA, MSB and MRI analyzed and interpreted experimental results as well as participated in manuscript preparations.

## Acknowledgement

The authors wish to thank Botanist Dr. Shaikh Bokhtear Uddin, Professor, Department of Botany, University of Chittagong, Bangladesh, who helped to identify the plant. We would like to express our gratitude to the authority of the International Centre for Diarrheal Disease and Research, Bangladesh (ICDDRB) for providing the experimental mice. The authors are also grateful to the Department of Pharmacy, International Islamic University Chittagong, Chittagong, Bangladesh, for providing research facilities.

## References

1. Chew AL, Jessica JJA, Sasidharan S. Antioxidant and antibacterial activity of different parts of *Leucas aspera*. Asian Pac J Trop Biomed. 2012; 2(3):176–180.

2. Buller R, Legrand V. Novel treatments for anxiety and depression: hurdles in bringing them to the market. Drug Discov Today. 2001; 6:1220–1230.

3. Yadav AV, Kawale LA, Nade VS. Effect of *Morus alba* L. (mulberry) leaves on anxiety in mice. Indian J Pharmacol. 2008; 40:32–36.

4. Titov N, Andrews G, Kemp A, Robinson E. Characteristics of adults with anxiety or depression treated at an internet clinic: comparison with a national survey and an outpatient clinic. PLOS ONE. 2010; 5(5):e10885.

5. Woode E, Abotsi WK, Mensah AY. Anxiolytic-and antidepressant-like effects of an ethanolic extract of the aerial parts of *Hilleria latifolia* (Lam.) H. Walt. in mice. J Nat Pharm. 2011; 2:62–71.

6. Katzung B: Basic and clinical pharmacology. 6th ed. California: Prentice-Hall International In. 1994, p. 323.

7. Rabbani M, Sajjadi SE, Mohammadi A. Evaluation of the anxiolytic effect of *Nepeta persica* Boiss. in mice. Evid-Based Complement Alternat Med. 2008; 5(2):181–186.

8. Rang HP, Dale MM, Ritter JM: Pharmacology. 3rd ed. London: Churchill Livingstone Inc.; 1996, p. 512.

9. Clark WG, Brater DC, Johnscn AR. Goth’s medical pharmacology. 12th ed. New Delhi: Galgotia Publication Pvt. Ltd. 1989, p. 288.

10. Essig CF. Addiction to nonbarbiturate sedative and tranquilizing drug. Clin Pharmacol Ther. 1964; 5:334–343.

11. Isbell H, Fraser HF. Addiction to analgesics and barbiturates. J Pharmacol Exp Ther. 1950; 99(4:2):355–397.

12. O’Brien CP. Drug addiction and drug abuse. In: Brunton L, Chabner B, Knollman B, editors. Goodman and Gilman’s the pharmacological basis of therapeutics. 9th ed. New York: McGraw-Hill Professional; 1996, p. 570.

13. Tripathi KD. Essential medical pharmacology. 3rd ed. New Delhi: Jaypee Brothers Medical Publishers; 1994, p. 324.

14. Schuckit MA. A low level of response to alcohol as a predictor of future alcoholism. Am J Psychiatry. 1994; 151:184–189.

15. Hook F. Flemingia stricta Roxb. Fl Ind 1832; 3:342.

16. http://www.legumes-online.net/ildis/aweb/td076/td_16024.htm.

17. Motaleb MA, Hossain MK, Alam MK, Mamun MA, Sultana M. Commonly used medicinal herbs and shrubs by traditional herbal practitioners Glimpses from Thanchi upazila of Bandarban, p.88–89.

18. Rahman MA, Uddin SB, Wilcock CC. Medicinal Plants used by Chakma Tribe in Hill Tracts Districts of Bangladesh. Indian Journal of Traditional Knowledge 2007; 6(3): 508–517.

19. Uddin SN. Traditional Uses of Ethnomedicinal Plants of the Chittagong Hill Tracts. Bangladesh National Herbarium, Bangladesh 2006. p 992.

20. Khisha T, Karim R, Chowdhury SR, Banoo R. Ethnomedical Studies of Chakma Communities of Chittagong Hill Tracts, Bangladesh. Bangladesh Pharmaceutical Journal 2012; 15(1):59–67.

21. Tiwari P, Kumar B, Kaur M, Kaur G, Kaur H. Phytochemical screening and extraction: a review. Int Pharm Sci. 2011;1:98–106.

22. Gupta BD, Dandiya PC, Gupta ML. A psychopharmacological analysis of behavior in rat. Jpn J Pharmacol. 1971; 21:293.

23. Takagi K, Watanabe M, Saito H. Studies on the spontaneous movement of animals by the hole cross test; effect of 2-dimethylaminoethanol and its acyl esters on the central nervous system. Jpn J Pharmacol. 1971; 21:797–810.

24. Ferrini R, Miragoli G, Taccardi B. Neuro-pharmacological studies on SB 5833, a new psychotherapeutic agent of the benzodiazepine class. Arzneim-Forsch (Drug Res) 1974; 24:2029–2032.

25. Lister RG. The use of a plus-maze to measure anxiety in the mouse. Psychopharmacology (Berl). 1987; 92:180–185.

26. Pellow S, File SE. Anxiolytic and anxiogenic drug effects on exploratrory activity in an elevated plus-maze: A novel test if anxiety in rat. Pharmacol Biochem Behav. 1986; 24:525–529.

27. Raquibul Hasan SM, Hossain MM, Akter R, Jamila M, Mazumder EHM, Rahman S. Sedative and anxiolytic effects of different fractions of the *Commelina benghalensis* Linn. Drug Discov Ther. 2009; 3:221–227.

28. Sen AK, Bose S, Dutta SK. Comparative evaluation of CNS depressant activity of the flavonoid fractions from the fresh leaves and flowers of *Ixora coccinea* Linn. J Pharma Sci Tech. 2011; 1(1):54–56.

29. Hossain MM, Biva IJ, Jahangir R, Vhuiyan MMI. Central nervous system depressant and analgesic activity of *Aphanamixis polystachya* (Wall.) parker leaf extract in mice. Afr J Pharm Pharmacol. 2009; 3(5):282–286.

30. Dhonnchadha BAN, Bourin M, Hascoet M. Anxiolytic-like effects of 5-HT2 ligands on three mouse models of anxiety. Behav Brain Res. 2011; 140:203–214.

31. Subramanian N, Jothimanivannan C, Kumar RS, Kameshwaran S. Evaluation of antianxiety activity of *Justicia gendarussa* burm. Pharmacologia. 2013; 4(5):404–407.

